# Structural evidence for two-stage binding of mitochondrial ferredoxin 2 to the core iron-sulfur cluster assembly complex

**DOI:** 10.1101/2024.02.19.580858

**Authors:** Ralf Steinhilper, Sven-A. Freibert, Susann Kaltwasser, Roland Lill, Bonnie J. Murphy

**Affiliations:** Redox and Metalloprotein Research Group, Max Planck Institute of Biophysics, Max-von-Laue-Str. 3, 60438 Frankfurt am Main, Germany; Institut für Zytobiologie, Philipps-Universität Marburg, Karl-von-Frisch-Str. 14, 35032 Marburg, Germany; Zentrum für Synthetische Mikrobiologie Synmikro, Karl-von-Frisch-Str. 14, 35032 Marburg, Germany; Central Electron Microscopy Facility, Max Planck Institute of Biophysics, Max-von-Laue-Str. 3, 60438 Frankfurt am Main, Germany

## Abstract

Iron-sulfur (FeS) clusters are ubiquitous metallocofactors that are essential for life. In eukaryotes, FeS cluster biosynthesis begins with the *de novo* assembly of a [2Fe-2S] cluster by the core iron-sulfur cluster assembly (ISC) complex in the mitochondrial matrix. This complex comprises the scaffold protein ISCU2, the cysteine desulfurase subcomplex NFS1-ISD11-ACP1, the allosteric activator frataxin (FXN) and the electron donor ferredoxin 2 (FDX2). The interaction of FDX2 with the complex remains unclear. Here, we present cryo-EM structures of the FDX2-bound core ISC complex and show that FDX2 and FXN compete for overlapping binding sites during [2Fe-2S] cluster biosynthesis. FDX2 binds in two conformations; in the ‘distal’ conformation, helix F of FDX2 shows loose electrostatic interaction with an arginine patch of NFS1, while in the ‘proximal’ conformation this interaction tightens and the FDX2-specific C terminus forms contacts with NFS1; in this conformation, the [2Fe-2S] cluster of FDX2 is close enough to the ISCU2 FeS cluster assembly site for rapid electron transfer.

## Introduction

Iron-sulfur (FeS) clusters are ancient metallocofactors that are essential for various cellular processes including oxidative phosphorylation, the citric acid cycle, and DNA replication and repair. In addition to their most prominent role in electron transfer and storage, FeS clusters can also be directly involved in catalysis or mediate structural support (*1, 2*). The biosynthesis of FeS clusters is carried out by protein-based machineries present in almost all living organisms (*3*). In eukaryotes, the process involves the mitochondrial iron-sulfur cluster assembly (ISC) machinery and the cytosolic iron-sulfur protein assembly (CIA) system, and catalyzes the biosynthesis of both [2Fe-2S] and [4Fe-4S] clusters in mitochondria, cytosol, and nucleus.

The process is initiated by the mitochondrial core ISC complex, which synthesizes a [2Fe-2S] cluster *de novo* from cysteine-derived sulfur and ferrous iron (Fe^2+^) (*4, 5*). This complex is composed of the iron-sulfur scaffold protein ISCU2, the dimeric pyridoxal phosphate (PLP)-dependent cysteine desulfurase subcomplex NFS1-ISD11-ACP1, the allosteric regulator frataxin (FXN), and the electron donor ferredoxin 2 (FDX2), which is itself reduced by the NADPH-dependent ferredoxin reductase (FDXR) (*6-11*). The reaction begins with the desulfuration of free L-cysteine at the PLP cofactor of NFS1 to produce L-alanine and a persulfide (-SSH) at residue Cys381^NFS1^ (*12*). The strictly conserved Cys381 is located on a flexible loop of NFS1 (termed ‘Cys-loop’) allowing the movement of the persulfide over a distance of more than 20 Å to the FeS cluster assembly site of ISCU2 which binds Fe^2+^ (*13, 14*). Persulfide transfer from Cys381^NFS1^ to Cys138^ISCU2^, one of three conserved cysteine residues of ISCU2, is facilitated by FXN, which was shown to increase persulfide transfer rates (*15-17*) by changing the local environment of the ISCU2 assembly site (*18, 19*). The sulfane sulfur (S^0^) of the persulfide on ISCU2 is reduced to sulfide (S^2-^) by FDX2 to initiate [2Fe-2S] cluster formation (*10*). Recent *in vitro* studies have shown that probably one electron is provided by FDX2, whereas a second electron could be derived from the oxidation of ISCU2-bound Fe^2+^ to ferric iron (Fe^3+^) (*14*). Tyr35^ISCU2^-induced dimerization of two ISCU2 proteins, assumed to each contribute a [1Fe-1S] moiety, is required to form a [2Fe-2S] cluster on ISCU2 (*20-22*). [2Fe-2S] units, generated by the core ISC complex, can be inserted into recipient proteins or serve as building blocks for the assembly of [4Fe-4S] clusters by components of the late ISC machinery (*4, 23*).

Mutations in genes encoding components of the core ISC complex are associated with iron accumulation in mitochondria and have been linked to several rare human diseases (*24*). Friedreich’s ataxia, the most common dysfunction of FeS cluster biosynthesis, is caused by decreased function of FXN in mitochondria, resulting in neuronal degeneration (*25*). Pathologies arising from mutations in the FDX2 and FDXR genes have been reported (*26-28*). Understanding the interplay of FXN and FDX2 with the core ISC complex on the molecular level will contribute to a fundamental understanding of the essential process of FeS cluster biosynthesis and may contribute to efforts at treating diseases caused by aberrant FeS cluster formation.

Despite several attempts to map ferredoxin binding to the core ISC complex, the precise interaction remains unknown. Competitive binding of bacterial frataxin (CyaY) and ferredoxin (Fdx) on the IscS dimer was proposed from nuclear magnetic resonance (NMR) spectroscopy (*29, 30*) and more recently from biochemical experiments using the yeast system (*31*) suggesting that the proteins bind in a sequential and transient fashion. In contrast, a model based on NMR data (recording the interaction of yeast ferredoxin (Yah1) with Isu1) and small-angle X-ray scattering (SAXS) envelopes of *Chaetomium thermophilum* homologues of the ISC machinery suggested a simultaneous binding of frataxin and ferredoxin to distinct sites (*13*). Humans express two ferredoxin isoforms, FDX1 (aka adrenodoxin) and FDX2 (*32*). FDX1 is most abundant in adrenal gland tissues, where it is involved in steroidogenesis, and was recently shown to be essential for the formation of the lipoyl and heme *a* cofactors in all cell types (*11*). FDX2 is ubiquitously expressed and is crucial for both [2Fe-2S] and [4Fe-4S] cluster biosynthesis (*8*). The C-terminal residues of the two human isoforms play a decisive role in defining their target specificity (*11*).

Previous X-ray crystallography and electron cryo-microscopy (cryo-EM) structures delivered important information on the overall architecture of the human core ISC complex (*13*) and its mode of interaction with FXN (*18*). However, these studies were performed in an aerobic atmosphere with copurified zinc (Zn^2+^) in the ISCU2 assembly site (*33*). The presence of Zn^2+^ is incompatible with the physiological [2Fe-2S] cluster formation because persulfide reduction by FDX2 is precluded in Zn^2+^-bound ISCU2 (*14*).

In this study, we present high-resolution cryo-EM structures of the human Fe^2+^-containing FDX2-bound core ISC complex by preparing grids under anaerobic conditions employing NADPH-FDXR-reduced FDX2. We show that FDX2 binding can occur in two related conformations, only one of which is likely to allow efficient electron transfer. Structural snapshots obtained by vitrification of turnover samples reveal that FDX2 and FXN compete for binding to overlapping sites at the core ISC complex during *de novo* [2Fe-2S] cluster biosynthesis.

## Results

### Structure determination of FDX2-bound core ISC complexes under anaerobic conditions

The initial aim of this study was to obtain cryo-EM structures of the biosynthetic NFS1-ISD11-ACP1-ISCU2 (hereafter abbreviated (NIAU)_2_) complex with bound FDX2 and FXN. To mimic near-physiological conditions for *de novo* FeS cluster biosynthesis during structure determination, we prepared and vitrified all cryo-EM samples under anaerobic conditions. FDX2 was reduced by catalytic amounts of FDXR and 1 mM NADPH in the buffer, because in this form yeast and human ferredoxins were shown to bind more tightly to the scaffold protein Isu1 (*10*). Cryo-EM samples were prepared under two different conditions (for details see Materials and Methods). First, to elucidate the interaction site of reduced FDX2 with the (NIAU)_2_ complex, the (NIA)_2_ subcomplex was incubated for 20 min with iron-loaded ISCU2 and FDXR-reduced FDX2 to yield the (NIAUF)_2_ sample. The second sample was prepared in exactly the same manner, but was incubated on ice and contained equimolar amounts of FXN and reduced FDX2 (hence abbreviated (NIAUXF)_2_). Cysteine was added to allow *de novo* FeS cluster synthesis (‘turnover’ conditions). The sample was quickly vitrified, resulting in a reaction time of less than 1 min at :< 4°C (*19*).

Structures of samples from both conditions were determined by single-particle cryo-EM, reaching resolutions of 2.0 Å for the (NIAUF)_2_ and 2.1 Å for the (NIAUXF)_2_ turnover conditions, respectively (Fig. S1 and Fig. S2). Until the consensus refinement step the overall architecture of the complexes appeared similar for the two conditions, showing a symmetric (NIAU)_2_ dimer with FDX2 tightly bound in a cavity between NFS1/NFS1’ and ISCU2 (Fig. 1). Additional weak density at this site (Fig. S3), typically associated with conformational and/or compositional heterogeneity in cryo-EM, was further assessed by C2 symmetry expansion and focused 3D classification (Fig. S4 and Fig. S5). This revealed two major FDX2 conformations that we refer to as the ‘proximal’ and ‘distal’ conformations according to the distances between the [2Fe-2S] cluster of FDX2 and the FeS cluster assembly site of ISCU2, namely 23 Å for the distal and 14 Å for the proximal conformation (Fig. 2A,B). In the (NIAUF)_2_ dataset, 61% of particles belong to a class in which FDX2 is bound in the proximal conformation, and 39% belong to a distal-bound FDX2 class (Fig. S4). For the turnover dataset, heterogeneity at the FDX2 binding site could be separated into three major classes (Fig. S5).

**Fig. 1.**
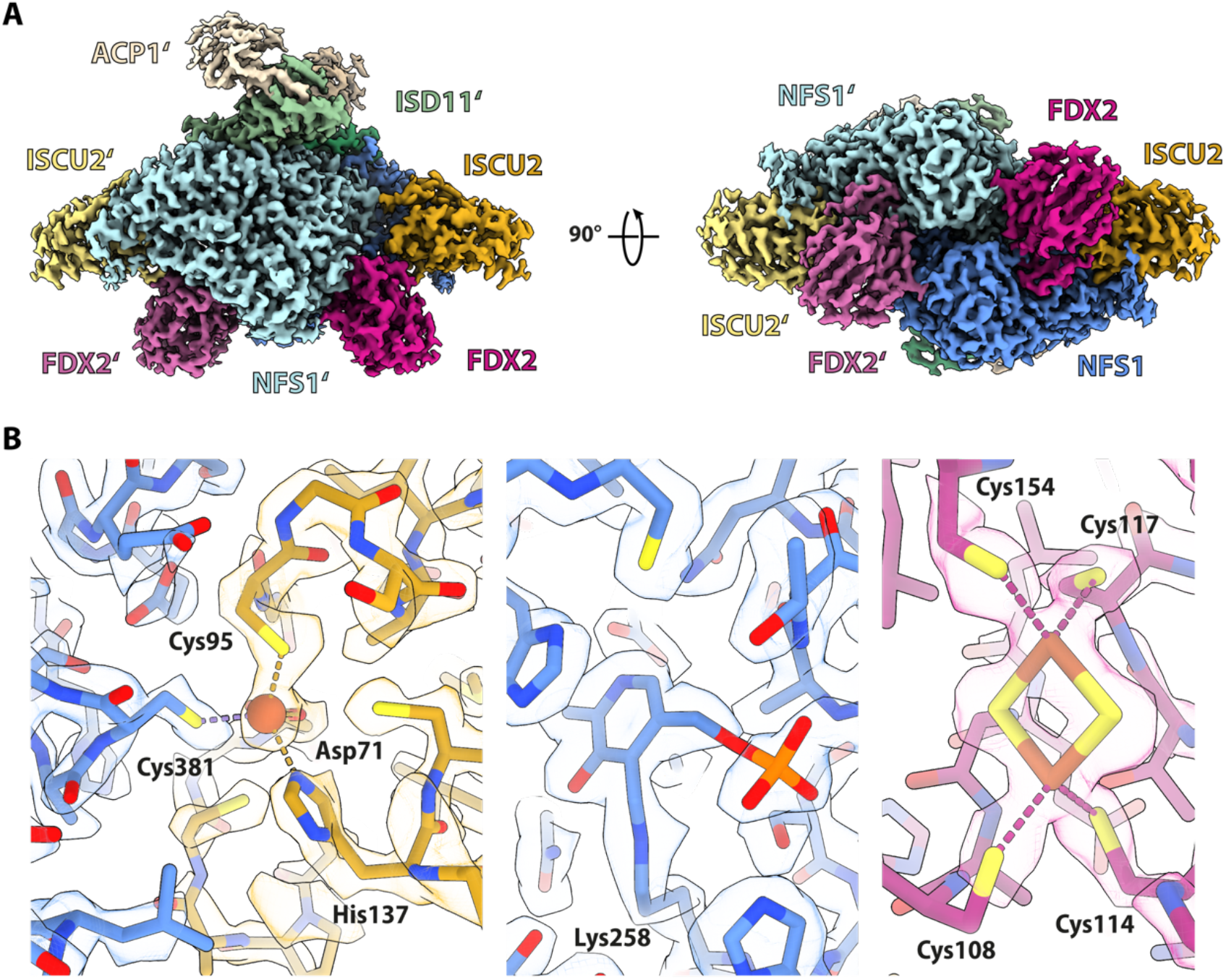
Overall architecture and cryo-EM density details of the (NIAUF)_2_ complex. (A) The 2 Å consensus cryo-EM map (C2 symmetry applied) is segmented and colored by subunit. Except where otherwise indicated, subunit coloring is consistent throughout the manuscript. FDX2 binds in a cavity at the NFS1-ISCU2 interface. (B) Atomic model and density details in the FDX2-bound proximal structure (density-modified). (Left) ISCU2 FeS assembly site with the Fe^2+^ ion (red sphere) coordinated by Cys381^NFS1^, Cys95^ISCU2^, Asp71^ISCU2^ and His137^ISCU2^; (middle) NFS1 PLP cofactor covalently linked to Lys258^NFS1^; (right) FDX2 [2Fe-2S] cluster coordinated by Cys108, Cys114, Cys117 and Cys154.

**Fig. 2.**
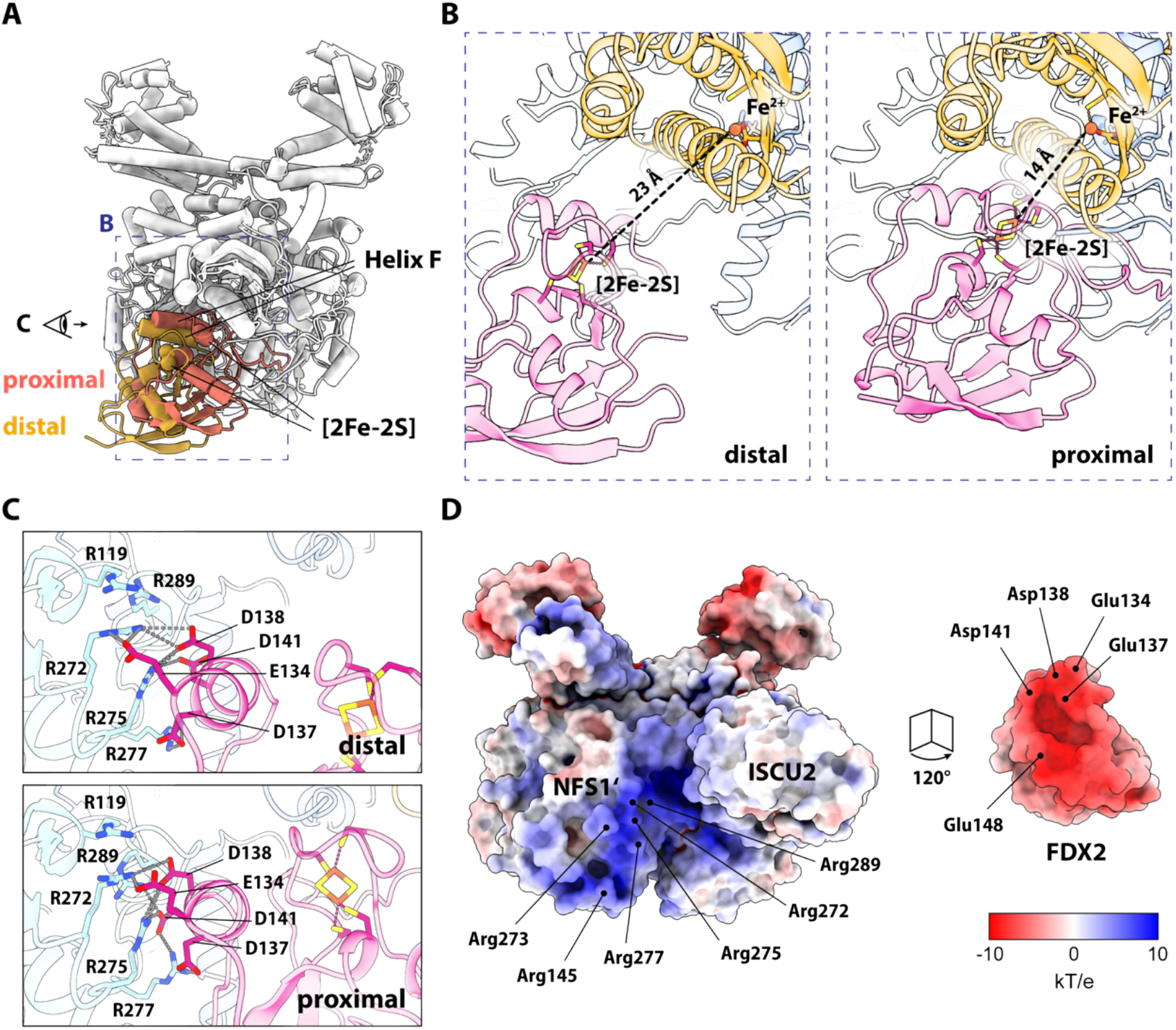
FDX2 binds the (NIAU)_2_ complex mainly via an arginine patch on NFS1’. (A) Overlay of the FDX2 distal (gold) and FDX2 proximal (orange) conformations. The conformational changes are visualized in Movie S1. Residues on helix F form salt bridges with NFS1’. The dashed blue box and the eye symbol represent the views shown in (B) and (C), respectively. (B) The distance between the FDX2 [2Fe-2S] cluster and the ISCU2 assembly site iron is 23 Å in the distal conformation. Binding of FDX2 in the proximal conformation decreases the distance to 14 Å. (C) Salt bridge interaction between FDX2 helix F and NFS1’ in the distal and proximal conformations. Distances <4 Å are indicated by dashed grey lines. (D) Electrostatic potential representation of the (NIAU)_2_ complex (left) and FDX2 (right). Residues involved in salt-bridge interactions between the patches are labelled.

The major proportion corresponds to FDX2 bound in the distal conformation (65%) and only 14% of the particles show FDX2 in the proximal conformation. The remaining 20% of the dataset corresponds to the FXN-bound core ISC complex (abbreviated (NIAUX)_2_), in which FXN is bound in a position virtually identical to that observed in previously published cryo-EM structures (*18, 19*).

### FDX2 interacts with a conserved arginine patch of NFS1

In both conformations (proximal and distal) FDX2 forms salt bridges with a conserved arginine patch (Arg272, Arg275, Arg277) and with Arg145, Arg273 and Arg289 of NFS1’ (NFS1 sequence alignment in Fig. S6). The contacts are made to acidic residues of FDX2 helix F (Glu134, Asp137, Asp138, Asp141) and Glu148^FDX2^, which interacts with Arg145^NFS1’^ (Fig. 2C,D). In the distal conformation, fewer salt bridges are formed with NFS1’ than in the proximal conformation (Fig. 2C) and the overall resolution of FDX2 is lower than for the rest of the complex (Fig. S1E), indicating a more flexible binding mode in the distal position than in the proximal conformation, where FDX2 appears well resolved (Fig. S1D).

The acidic patch of FDX2 helix F is highly conserved (FDX2 sequence alignment in Fig. S7) and was previously suggested as the binding interface with NFS1 based on NMR experiments using the bacterial homologues (*29, 30*). This region additionally acts as the primary binding interface with the ferredoxin reductase (*34*). The NFS1 arginine patch has also been previously proposed to serve as a nuclear localization signal (*35*), although we note that the motif is also present in bacterial sequences (Fig. S6).

### Binding of FDX2 in the proximal conformation stabilizes the flexible C terminus and brings the [2Fe-2S] cluster in a position suited for efficient electron transfer

In the distal conformation, the [2Fe-2S] cluster of FDX2 is too far for efficient electron transfer to the ISCU2 FeS cluster assembly site (23 Å) (*36*) (Fig. 2B), whereas the movement to the proximal binding position brings the [2Fe-2S] cluster to a distance of 14 Å from the FeS cluster assembly site, a distance that would allow rapid electron transfer for reduction of the Cys138^ISCU2^ persulfide (Fig. 2B). In the distal binding position, the C-terminal 15 residues of FDX2 are unresolved, indicating that these residues are highly flexible when FDX2 is bound in this conformation. Disordered C termini are common among ferredoxins, and this region had to be truncated in human FDX2 to facilitate its crystallization (PDB 2Y5C) (*11*). When bound in the proximal binding site, the C terminus of FDX2 is ordered and well resolved interacting extensively with the NFS1 subunit (Fig. 3A). Contact is mediated by hydrogen bonds (H-bonds) between Asn175^FDX2^ and Cys-loop Ser385^NFS1^ (Fig. 3B), a salt bridge between Asp179^FDX2^ and Arg393^NFS1^ (Fig. 3C) and hydrophobic interactions between Phe176^FDX2^ and Val178^FDX2^ with Leu160^NFS1^. The interacting residues are conserved among higher eukaryotes (sequence alignments in Fig. S6 and Fig. S7), which underlines the functional importance of these interactions. Unexpectedly, the highly conserved terminal Lys-Pro-His motif and other conserved C-terminal residues of FDX2 do not seem to make specific contacts with NFS1 or ISCU2 subunits. Therefore, the function of this well-conserved region may not solely be in the recognition of the core ISC complex. Rather, the C terminus may also play a role in other parts of the FeS protein biogenesis pathway, for instance during the fusion of two [2Fe-2S] to a [4Fe-4S] cluster by the ISCA1-ISCA2-IBA57 complex (*23*).

**Fig. 3.**
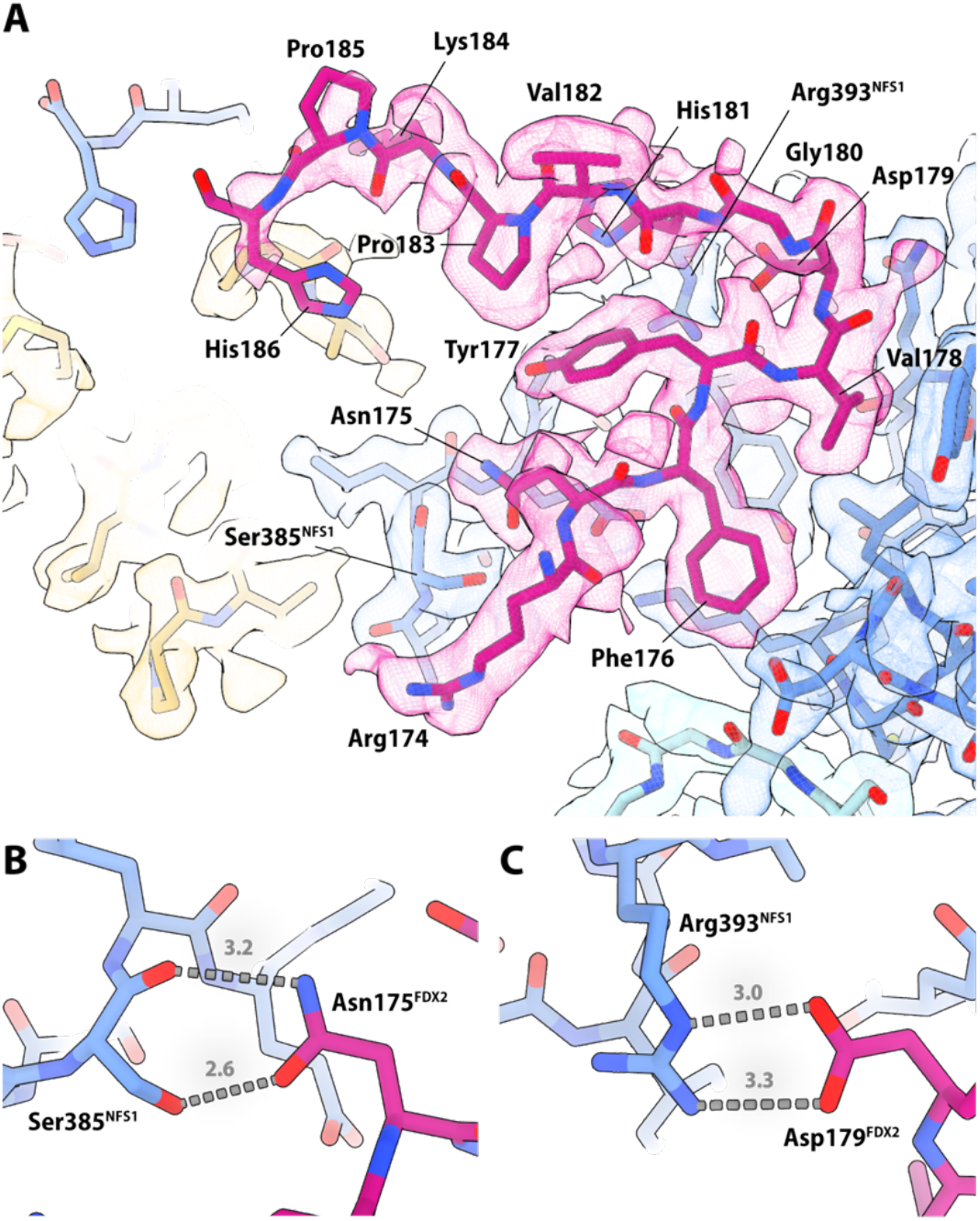
Density and atomic model of the FDX2 C terminus. (A) In the proximal conformation of the (NIAUF)_2_ structure the FDX2 C terminus is resolved and shows side-chain density for nearly all residues, except for Lys184, Pro185 and His186. (Only FDX2 residues 174-186 are displayed for figure clarity). (B) H-bonds can form between Asn175^FDX2^ of the FDX2 C terminus and Ser385^NFS1^ of the NFS1 Cys-loop. (C) Salt-bridge interaction occurs between Asp179^FDX2^ and Arg393^NFS1^.

The loop region of ISCU2 including Cys69^ISCU2^ rearranges in correlation with the two FDX2 conformations; when FDX2 is bound in the distal conformation, Cys69^ISCU2^ is shifted away from the ISCU2 FeS cluster assembly site (Fig. 4A, Movie S1), whereas it is facing towards the iron, but too far for coordination (3.1 Å) when FDX2 is bound in the proximal conformation (Fig. 4B). The functional implication of this rearrangement remains to be clarified.

**Fig. 4.**
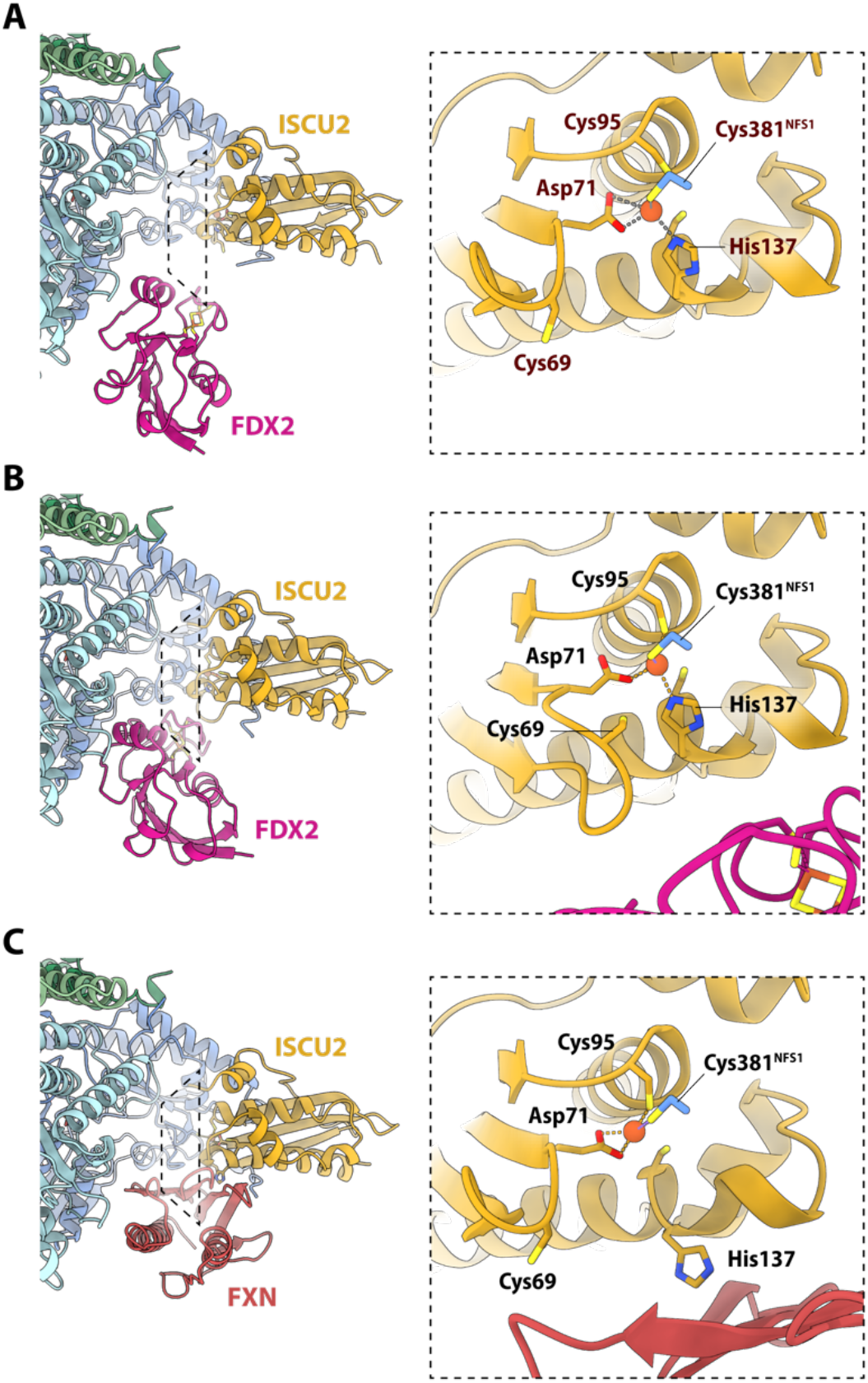
Iron coordination in the ISCU2 FeS cluster assembly site in the presence of FDX2 or FXN. (A) When FDX2 is bound in the distal conformation, the iron is coordinated by Asp71^ISCU2^, His137^ISCU2^ and Cys381^NFS1^. Cys95^ISCU2^ faces the iron, but is too far for coordination (3.5 Å). Cys69^ISCU2^ faces away from the ISCU2 FeS assembly site, similar to the FXN-bound structure. (B) When FDX2 is bound in the proximal conformation, the iron is coordinated by Asp71^ISCU2^, Cys95^ISCU2^, His137^ISCU2^ and Cys381^NFS1^. Cys69^ISCU2^ faces towards the iron, but is too far for coordination (3.9 Å). (C) In the FXN-bound structure, the iron in the ISCU2 FeS assembly site is coordinated by Asp71^ISCU2^, Cys95^ISCU2^ and Cys381^NFS1^.

### Ferredoxin and frataxin bind to overlapping binding sites during turnover

The conserved arginine residues of NFS1’ (Arg272, Arg275, Arg277, Arg289), which electrostatically interact with FDX2, also serve as the binding site for FXN (primarily via its residues Glu108, Glu111, Glu121 and Asp124) (*18, 37*). It was previously suggested that ferredoxin and frataxin compete for binding in homologous bacterial and yeast complexes (*29-31*). In the bacterial system, Kim *et al*. (*30*) found that ferredoxin, IscU and CyaY (FXN homolog) compete for an overlapping binding site on IscS (NFS1 homolog). This does not appear to be the case for the human core ISC complex, as our cryo-EM structures show ISCU2 being tightly bound in the presence of FDX2 or FXN.

In our FXN-bound (NIAUX)_2_ structure, the Cys-loop of NFS1 is predominantly facing the ISCU2 assembly site with Cys381^NFS1^ coordinating the iron (Fig. 4C), but additional weak density in this region may originate from a low-occupancy alternate conformation of the Cys-loop, consistent with its high flexibility. FXN interacts with His137^ISCU2^ and Cys69^ISCU2^, which was previously observed for the Zn^2+^ and Fe^2+^-bound (NIAUX)_2_ structures (*18, 19*) and likely modulates the ISCU2 assembly site to facilitate persulfide transfer to Cys138^ISCU2^. In contrast, the structures of both FDX2 conformations show His137^ISCU2^ facing the ISCU2 FeS cluster assembly site in coordinating distance to Fe^2+^. This conformation of His137^ISCU2^ is more similar to the structures of non-FXN-bound (NIAU)_2_ complexes.

All our FDX2-bound structures show the Cys-loop in the ‘outward’ conformation where Cys381^NFS1^ coordinates the ISCU2-bound Fe^2+^ together with Cys95^ISCU2^, Asp71^ISCU2^ and His137^ISCU2^ (Fig. 4A,B). When FDX2 is bound in the proximal position, the H-bonds that form between Asn175^FDX2^ and Ser385^NFS1^ could provide additional stability for the ‘outward’ Cys-loop conformation. An outward-facing conformation of the Cys-loop was also observed in the X-ray structure of Zn^2+^-bound (NIAU)_2_ (*13*), indicating that this may be a resting conformation when neither FXN nor FDX2 are bound to the (NIAU)_2_ complex.

## Discussion

Our cryo-EM structures reveal the binding sites of FDX2 to the human core ISC complex at near-atomic resolution. FDX2 interacts electrostatically with a series of arginine residues on NFS1’, overlapping with the binding position of FXN. This finding is consistent with previous studies that observed competitive binding of the bacterial FDX2 and FXN orthologs (*29, 30*). We did not observe any evidence for a dodecameric core ISC complex, in which FDX2 and FXN would bind simultaneously, as suggested by a previous model (*13*). This model was established on the basis of SAXS envelopes obtained from various reconstitutions using components of the *C. thermophilum* core ISC complex (*13*) taking into consideration interacting residues between *C. thermophilum* apo-Isu1 (ISCU2 homolog) and chemically reduced yeast Yah1 (FDX2 homolog) that were identified by NMR spectroscopy (*10*). In this model, FDX2 is suggested to interact with ISCU2 in a location that is occupied by the NFS1 C terminus in the cryo-EM structures presented here and by Fox *et al*. (*18*). In our present study, the tight and specific interaction between NFS1 and FDX2 via salt bridges and the rigidification of the otherwise flexible C-terminal residues supports the idea that this is the physiological binding site of FDX2 within the core ISC complex during FeS cluster biogenesis (Fig. 5).

**Fig. 5.**
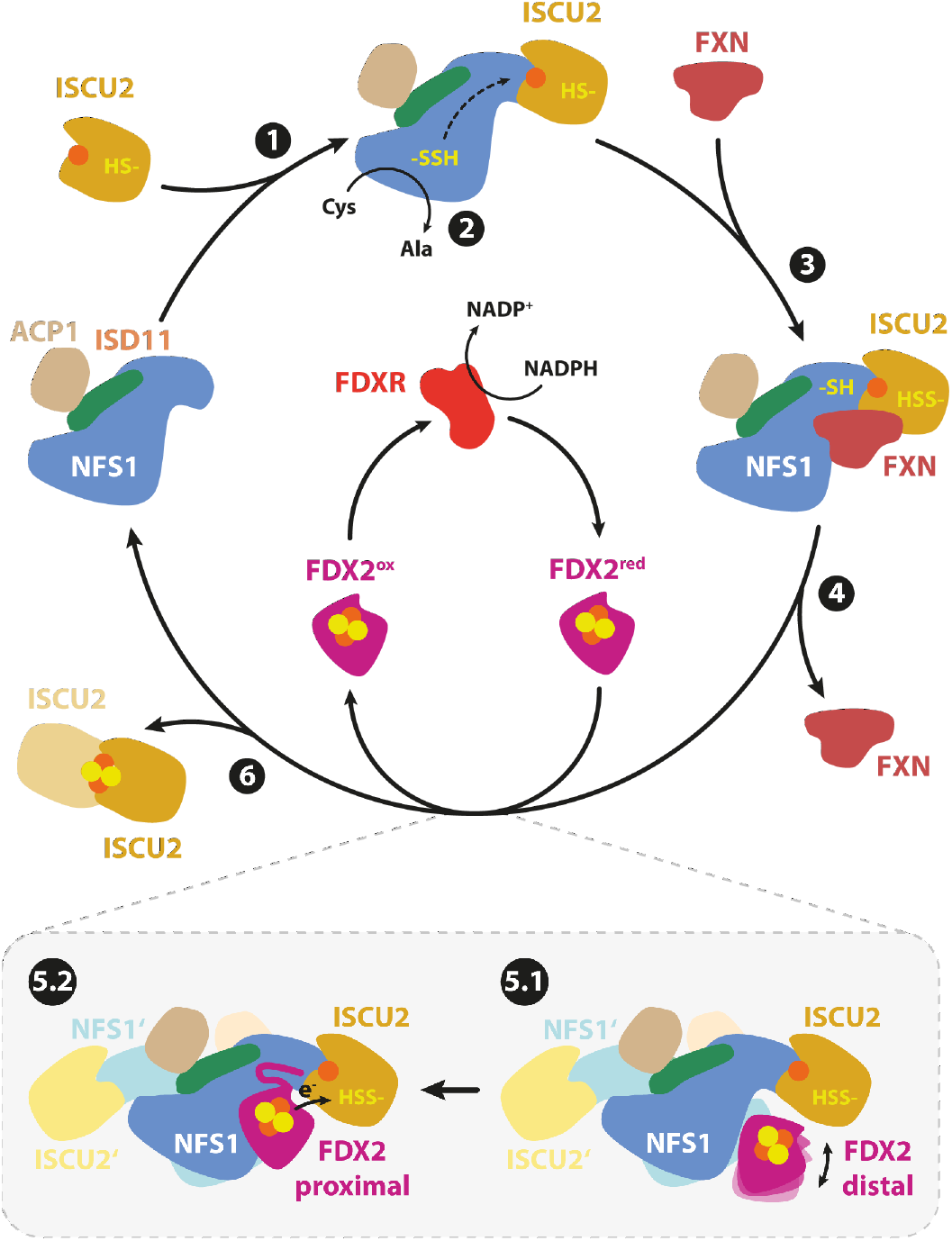
Scheme of the major steps of *de novo* [2Fe-2S] cluster biogenesis by the core ISC complex. For simplicity the second dimer-half is shown only for step 5. (1) The (NIA)_2_ subcomplex binds to the FeS scaffold protein ISCU2 to form the (NIAU)_2_ complex. (2) Desulfuration of free cysteine (Cys) at the PLP cofactor yields a protein-bound persulfide, which is transferred to ISCU2 via Cys381^NFS1^ of the flexible Cys-loop. (3) Transient interaction with the allosteric activator FXN modulates the ISCU2 assembly site to facilitate sulfur transfer from Cys381^NFS1^ to Cys138^ISCU2^. (4) FXN leaves the complex to allow FDX2-binding. (5.1) Reduced FDX2 binds loosely to the NFS1 arginine patch in the distal conformation. (5.2) Rigidification of the flexible FDX2 C terminus allows FDX2 to shift into the more rigidly bound proximal conformation where the [2Fe-2S] cluster is close enough for donation of an electron (e^-^) to the ISCU2 FeS cluster assembly site. Oxidized FDX2 is reduced by FDXR and NADPH. (6) The [1Fe-1S] intermediate is converted into a mature [2Fe-2S] cluster by dimerization of two ISCU2 subunits.

Our structures show that FDX2 binds in two conformations, termed ‘proximal’ and ‘distal’ according to the distance of the electron-donating [2Fe-2S] cluster of FDX2 to the ISCU2 FeS cluster assembly site, where Fe^2+^ and the persulfide bind. During the transition from the distal to the proximal conformation, the FDX2 [2Fe-2S] cluster moves by a distance of around 8 Å, thereby diminishing the distance to the ISCU2 FeS cluster assembly site from 23 to 14 Å, a distance that would allow efficient electron transfer. In the proximal conformation the conserved C-terminal residues of FDX2 become ordered and interact extensively with NFS1. The two conformations are related by a rotational motion around FDX2 helix F (Movie S1), which contains negatively charged residues that form electrostatic interactions with several arginine residues of NFS1. This movement involves a tightening of the salt bridges between NFS1’ and FDX2. A possible explanation for the existence of two binding modes for FDX2 could be that the distal conformation is kinetically favored and represents a first stage of binding, which then enables the intrinsically disordered C terminus of FDX2 to rigidify and bind to NFS1. Strong conservation of the ‘type II ferredoxin’ (FDX2) C-terminal residues is the key difference that distinguishes FDX2 from type-I adrenodoxin (FDX1) and the fungal-type eukaryotic [2Fe-2S] ferredoxins (*11, 32*). FDX2 plays a central role in FeS cluster biosynthesis (*8*) and its specificity partially depends on the C-terminal 12 residues (*11*). In this context, a likely role of the C terminus of FDX2 might therefore be to mediate the transition to a productive, electron-transfer competent complex. Nevertheless, not all of the conserved C-terminal residues make specific contacts to NFS1, suggesting additional functions of this segment in other steps of the biosynthetic process.

Alternately, it may be that the two modes of binding offer some measure of temporal control. Because FDX2 binding in the proximal position involves interaction with residues of the Cys-loop, it is conceivable that progression through the catalytic steps in the (NIAU)_2_ complex could shift the equilibrium between proximal and distal binding modes. Such a mechanism could be important in modulating the relative likelihood of FXN vs. FDX2 binding at different steps along the catalytic pathway. Further experiments would be required to assess these possibilities.

Overall, our structures fit to the model derived from biochemical studies that *de novo* [2Fe-2S] cluster biosynthesis by the core ISC complex is a highly concerted process with FXN and FDX2 binding sequentially rather than simultaneously (*31*) (Fig. 5). This causes distinct conformational changes in the ISCU2 FeS assembly site, which is activated by FXN binding to facilitate sulfur transfer from Cys381^NFS1^ to Cys138^ISCU2^, but is inactive in the FDX2-bound structures, similar to the situation observed when neither FDX2 nor FXN are bound to the complex. The fact that FDX2 and FXN compete for a binding site may have implications for understanding the concentration-dependent effects of FXN expression in Friedreich’s ataxia.

## Materials and Methods

### Protein production and purification

All proteins (sequence information listed in Table S1) were synthesized recombinantly in *Escherichia coli* strain BL21 (DE3) and purified at 4°C or on ice (*22*). FDXR was produced and purified as described previously (*23*). For all other proteins, cells were transformed with the appropriate plasmids (Table S2) and grown at 37°C for 8 h in terrific broth media. For FDX2 or (NIA)_2_ expression, *E. coli* media were supplemented with 50 μM Fe(NH_4_)_2_(SO_4_)_2_ or 2 mM pyridoxine hydrochloride, respectively. Gene expression was induced at OD_600_ ∼0.8 by addition of either 1 mM isopropyl-β-D-thiogalactopyranoside (IPTG; Carl Roth GmbH) or 0.2 µM anhydrotetracycline (Sigma). Cells were grown overnight at 22°C, harvested by centrifugation, and flash-frozen in liquid nitrogen. For purification of His-tagged proteins ((NIA)_2_, ISCU2 and FXN), cells were thawed at room temperature and resuspended in IMAC buffer (35 mM Tris-HCl pH 7.4, 300 mM NaCl, 5% (w/v) glycerol and 10 mM imidazole). FDX2 was purified by anion exchange chromatography (AEC), for which cells were resuspended in AEC buffer (35 mM Tris-HCl pH 7.5, 50 mM NaCl, 5% (w/v) glycerol). Protease inhibitor (cOmplete; Roche), lysozyme, and DNase I (and 5 mM PLP for (NIA)_2_ purification) were added to the cell suspensions prior to cell lysis by sonication (SONOPULS mini20; BANDELIN electronic GmbH & Co. KG). Lysates were cleared by centrifugation at 40,000 xg for 45 min. Supernatants of (NIA)_2_, ISCU2 and FXN were subjected to Ni-NTA affinity chromatography (His-Trap 5ml FF crude; GE Healthcare). The column was washed with 10 CV IMAC wash buffer, which contained 10 mM imidazole (for (NIA)_2_) or 70 mM imidazole (for ISCU2 and FXN). Proteins were eluted with IMAC buffer containing 250 mM imidazole.

The supernatant of FDX2 was loaded onto an anion exchange column (Source 30Q) pre-equilibrated with AEC buffer and eluted with a linear gradient to high-salt buffer (35 mM Tris-HCl pH 7.5, 1 M NaCl, 5% (w/v) glycerol). Eluted proteins were concentrated. For (NIA)_2_ purification an additional AEC step was added after His purification to achieve higher purity for structural studies. FXN was treated with recombinant TEV protease (purified in-house) for removal of the N-terminal His-tag. Proteins were transferred into an anerobic chamber (Coy Laboratories) and incubated for 60 min with 10 mM DTPA, 10 mM TCEP, 25 mM KCN (for FXN and ISCU2) and 0.5 mM PLP (additionally for (NIA)_2_) to remove metal ions and polysulfane sulfur before proteins were subjected to anaerobic size exclusion chromatography (SEC) (HiLoad 16/600 Superdex; GE Healthcare) on an Äkta Purifier system (GE Healthcare) using SEC buffer (50 mM Tris-HCl pH 7.4, 150 mM NaCl, 5% glycerol). ISCU2 was pre-incubated with FeCl_2_ and ascorbate (both in 5x molar excess), FDXR was supplemented with MgCl_2_ in 2x molar excess. Buffer was exchanged to 20 mM Tris-HCl pH 7.4, 100 mM NaCl for all proteins with PD10 columns (GE Healthcare) and protein concentrations were determined by the Bradford assay (Biorad). A correction factor for the resulting concentrations was determined for (NIA)_2_, ISCU2, FXN, and FDX2 by quantitative amino acid analysis (Leibniz-Institut für Analytische Wissenschaften). Proteins were transferred into gas-tight glass vials and flash-frozen in liquid N_2_.

### Cryo-EM sample preparation

Samples for single-particle cryo-EM analysis were prepared under anaerobic conditions in an atmosphere of 3-5% H_2_ in an N_2_ atmosphere within an anaerobic tent (Coy Laboratories). Protein stock solutions were thawed on ice and anaerobic cryo-EM buffer (20 mM Tris-HCl pH 7.4, 100 mM NaCl, 1.1 mM NADPH) was prepared fresh. For the (NIAUF)_2_ sample,

27.5 µM (NIA)_2_ was mixed with 110 µM iron-loaded ISCU2 to yield the (NIAU)_2_ complex. Subsequently, 110 µM holo-FDX2 and catalytic amounts (2.75 µM) of FDXR were added and the sample was incubated for 20 min at room temperature in cryo-EM buffer. 2.7 µL of sample was mixed with 0.3 µL of fluorinated fos-choline-8 solution (Anatrace; final concentration 1.5 mM) and applied to UltrAuFoil 0.6/1 300 mesh gold grids (Quantifoil), which had been glow-discharged two times for 90 s at 15 mA and 0,38 mbar using a PELCO easiGlow unit (Ted Pella Inc.). The sample was blotted at 4°C and 100 % humidity and vitrified in liquid ethane using a Vitrobot Mark IV (Thermo Scientific). The (NIAUXF)_2_ turnover sample was prepared similarly, but 110 µM FXN and 300 µM cysteine were added, prior to vitrification, resulting in a total reaction time of less than 1 min at :< 4°C. We used gold support grids for our study because of their inert character and their advantage in limiting beam-induced motion during data acquisition (*38*).

### Data acquisition and image processing

Cryo-EM data were collected at 300 kV on a Krios G4 (Thermo Scientific) equipped with a cold field emission gun (E-CFEG) and a Falcon 4 direct electron detector with a Selectris X Imaging filter. A nominal magnification of 215,000x was used, which corresponds to a calibrated pixel size of 0.573 Å. For the (NIAUF)_2_ sample, a dataset of 8497 EER files (952 internal frames) was collected in counting mode with a total dose of 80 e^-^/Å^2^ using aberration-free image shift (AFIS) in EPU (Thermo Scientific) (Table S3). For the (NIAUXF)_2_ turnover sample, two datasets were collected from the same grid with identical settings (Table S4); dataset 1 contained 9144 EER files (987 internal frames), dataset 2 contained 8747 EER files (1078 internal frames). These datasets were processed separately until the particle polishing step (Fig. S5).

The overall data processing strategy until particle polishing of the three datasets was similar (Fig. S4 and Fig. S5) and will hereafter be exemplarily elucidated for the (NIAUF)_2_ dataset. Movies were sorted into optics groups corresponding to their EPU AFIS metadata (https://github.com/DustinMorado/EPU_group_AFIS) before they were gain-normalized and motion corrected with dose-weighting using MotionCor2 (*39*) in RELION-4 (*40*). The contrast transfer function (CTF) was estimated with CTFFind4.1.13 (*41*). Particle coordinates, initially picked with the blob picker implementation in cryoSPARC live (*42*) during data acquisition of the respective datasets and cleaned by 2D classification, were imported into RELION and used to train a particle detection model with Topaz (*43*). Topaz picking parameters were first tested and adjusted on a small particle subset before particles were picked from all micrographs and extracted using a box size of 416x416, rescaled to 104x104 (pixel size 2.292 Å). These particles were then imported to cryoSPARC v4.1.2 (*42*) and subjected to 2D classification. Three ab-initio 3D reconstructions were generated from select particles, and used as templates for heterogenous refinement (3 classes, no symmetry applied). The major class (510,575 particles), revealing the characteristic appearance of the core ISC complex, was selected and subjected to homogenous refinement (no symmetry applied). The refined particle coordinates were converted into a particle STAR file using pyem (*44*) and imported into RELION for re-extraction using a box size of 416x416 rescaled to 288x288 (pixel size 0.828 Å), and imported to cryoSPARC for a non-uniform refinement applying C2 symmetry and optimizing per-group defocus and CTF parameters, which yielded a reconstruction at a resolution of 2.33 Å. Finally, these refined particles were imported into RELION for Bayesian particle polishing using custom parameters (trained on 10,000 particles). The polished particles were imported to cryoSPARC and subjected to a non-uniform refinement (C2 symmetry applied, per-group defocus and CTF optimization enabled), followed by a heterogenous refinement (2 classes, C2 symmetry applied). Particles from the major 3D class (363,652 particles) were subjected to a non-uniform refinement (C2 symmetry applied, per-group defocus and CTF optimization enabled), resulting in a consensus map with a global resolution of 2.03 Å. For the (NIAUXF)_2_ turnover datasets, the global resolutions for the consensus non-uniform refinement of the two combined datasets gave a global resolution of 2.09 Å (C2 symmetry applied, per group defocus and CTF optimization enabled). The overall appearance of the two consensus maps was highly similar, but some degree of heterogeneity could be observed at the FDX2 binding site (Fig. S1C, Fig. S2C and Fig. S3), which was further investigated by performing C2 symmetry expansion and focused 3D classification.

Particles from the (NIAUF)_2_ consensus refinement were C2 symmetry expanded with *relion_particle_symmetry_expand* and subjected to a focused 3D classification without alignment (number of classes = 2, regularisation parameter T=4, number of iterations = 50, initial low-pass filter = 10 Å) using a mask on the FDX2-binding region. This separated the particles into two classes; Class 1 (38.8%) shows FDX2 bound to NFS1 in the distal conformation, whereas Class2 (61.2%) shows FDX2 bound in the groove between NFS1 and ISCU2, in the proximal conformation. Particles from both classes were imported into cryoSPARC for local refinements (no symmetry applied) yielding resolutions of 2.39 Å (Class 1) and 2.26 Å (Class 2; FDX2-bound proximal). Particles from Class 1 were imported into RELION for particle subtraction using a mask around the FDX2-binding region and subjected to a focused 3D classification without alignment (number of classes = 4, regularisation parameter T=256, number of iterations = 50, initial low-pass filter = 10 Å). 88,788 particles from the major class (Class 3; 31.1%) were reverted to the original particles, imported to cryoSPARC and subjected to a local refinement (no symmetry applied) yielding a resolution of 2.52 Å (FDX2-bound distal).

For the (NIAUXF)_2_ turnover dataset, polished particles from both datasets were subjected to a non-uniform refinement (C2 symmetry applied, per group defocus and CTF optimization enabled), which converged at a resolution of 2.09 Å. The 731,041 particles from this consensus refinement were converted into a STAR file using pyem, C2 symmetry expanded with *relion_particle_symmetry_expand* and subjected to a focused 3D classification without alignment (number of classes = 3, regularisation parameter T=16, number of iterations = 50, initial low-pass filter = 10 Å) using a mask around the FDX2-binding region. This separated the dataset into classes with FXN bound (20.4%), FDX2 bound in the proximal conformation (14.2%) and FDX2 bound in the distal conformation (65.4%). Particles from each class were imported into cryoSPARC and subjected to local refinements, reaching resolutions of 2.49 Å (FXN-bound), 2.33 Å (FDX2-bound proximal) and 2.35 Å (FDX2-bound distal). Particles of the FDX2-bound distal class were imported to RELION for particle subtraction using a mask on the FDX2-binding region before they were subjected to a focused 3D classification without alignment (number of classes = 4, regularisation parameter T=128, number of iterations = 50, initial low-pass filter = 10 Å). The 3D class with the strongest density was selected (326,204 particles), reverted to the original particle images, imported into cryoSPARC and subjected to a local refinement (no symmetry applied), reaching a resolution of 2.46 Å.

### Model building and refinement

Half maps of the ‘FDX2-bound proximal’ and ‘FDX2-bound distal’ structures originating from the (NIAUF)_2_ dataset were used for density modification with *phenix*.*resolve_cryo_em* (*45*) (molecular mass provided, ‘real-space weighting’ and ‘weight by sigmas’ options enabled). This improved the overall map interpretability and facilitated model building and identification of water molecules. For model refinement of the structures of the (NIAUXF)_2_ turnover dataset, the auto-sharpened maps were used, except for the ‘FXN-bound’ structure, which was manually sharpened (B-factor = -60 Å^2^).

Structures of the human Zn^2+^-bound (Zn-NIAUX)_2_ complex (PDB 6NZU) (*18*) and human FDX2 (PDB 2Y5C) (*10*) served as templates for the ‘FDX2-bound proximal’ and ‘FDX2-bound distal’ structures and were rigid-body fitted into the respective cryo-EM maps using ChimeraX (*46*). The coordinates were real-space refined against the respective maps using *phenix*.*real_space_refine* within PHENIX (*47*) followed by manual fitting in Coot-0.9 (*48*). Metal restraints were added with *ReadySet!* in PHENIX, and adjusted manually according to published high-resolution structures. Waters were built automatically with *phenix*.*douse* and checked manually for sensible hydrogen bonding. Final models were validated by *phenix*.*validation_cryoem*.

### Visualization

Sequence alignments were performed with ClustalW (*49*) and visualized with the ESPript server (*50*). All maps and models were visualized in ChimeraX (*46*). The electrostatic potential map in Fig. 2D was calculated with the APBS-PDB2PQR software suite (*51*) and displayed in ChimeraX.

## Acknowledgments

We thank the Central Electron Microscopy Facility of the Max Planck Institute of Biophysics for providing cryo-EM infrastructure and technical support, and Werner Kühlbrandt and the Department of Structural Biology for support and helpful discussions. We acknowledge the contribution of the Core Facility ‘Protein Biochemistry and Spectroscopy’ of the Philipps-Universität Marburg.

## Author contributions

RS prepared cryo-EM samples, acquired and processed cryo-EM data, built and refined the atomic models, analyzed data, drew the figures and wrote the initial draft. SAF purified the proteins and revised the manuscript. SK supported cryo-EM data acquisition and revised the manuscript. RL and BJM initiated the study and supervised the project, analyzed data, wrote and edited the manuscript.

## Data and materials availability

Cryo-EM maps and atomic models from the (NIAUF)_2_ dataset were deposited to the Electron Microscopy Data Bank and the Protein Data Bank under the accession codes EMD-19355 ((NIAUF)_2_ consensus map), EMD-19356 and PDB 8RMC (FDX2-bound proximal), EMD-19357 and PDB 8RMD (FDX2-bound distal). Cryo-EM maps and atomic models originating from the (NIAUXF)_2_ turnover datasets are accessible under EMD-19358 ((NIAUXF)_2_ turnover, consensus map), EMD-19359 and PDB 8RME ((NIAUXF)_2_ turnover, FXN-bound), EMD-19360 and PDB 8RMF ((NIAUXF)_2_ turnover, FDX2-bound proximal), EMD-19361 and PDB 8RMG ((NIAUXF)_2_ turnover, FDX2-bound distal). Raw data are available from the corresponding authors upon request.

