## Supplementary Materials for "Structural evidence for two-stage binding of mitochondrial ferredoxin 2 to the core iron-sulfur cluster assembly complex"

**This PDF file includes:**

Figs. S1 to S7
Tables S1 to S4
Captions for Movie S1
References

**Other Supplementary Materials for this manuscript include the following:**
Movie S1

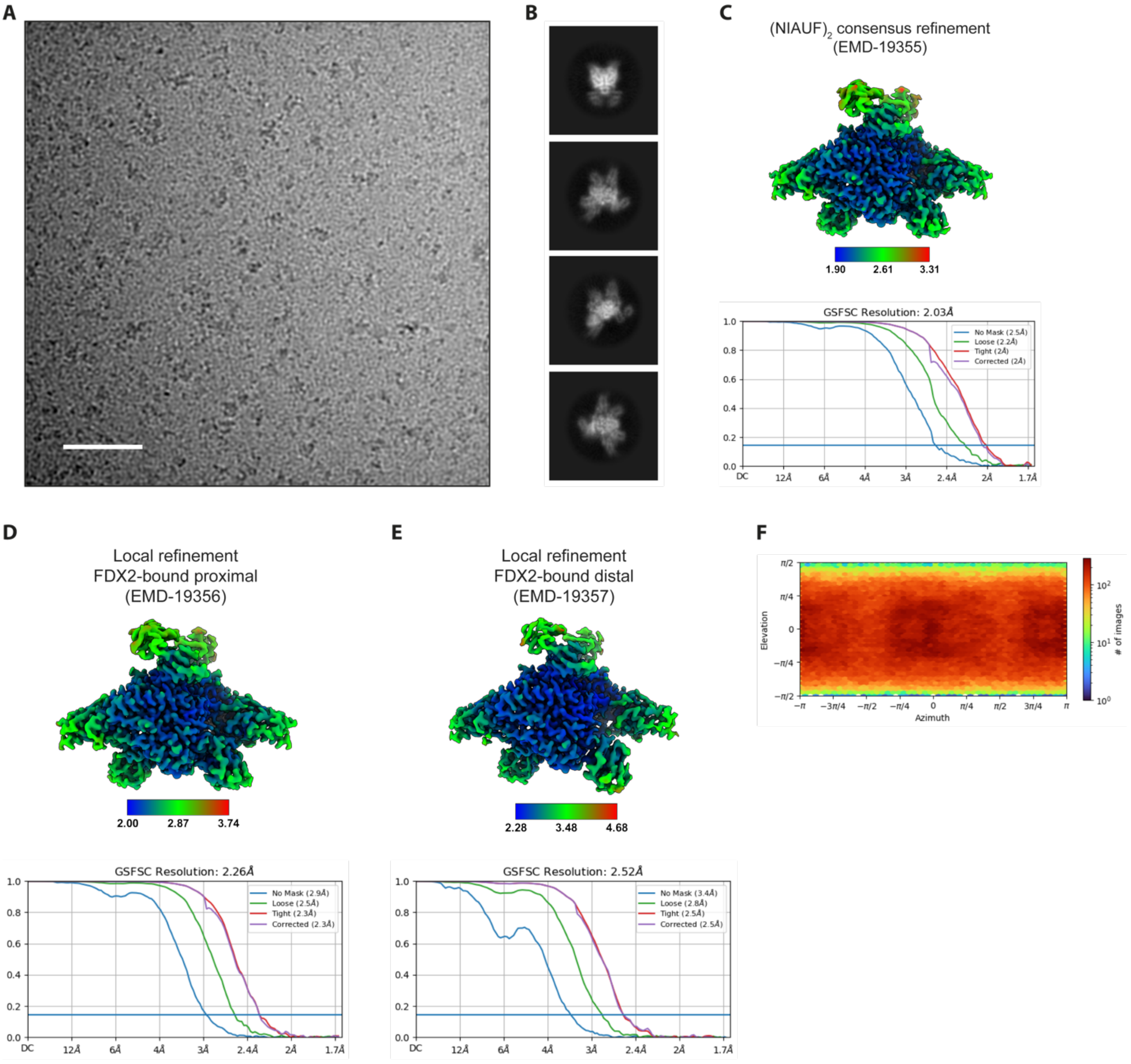


**Fig. S1. Cryo-EM data processing of the $(NIAUF)_2$ dataset.**

(**A**) Representative cryo-EM micrograph (scale bar 40 nm), (**B**) representative 2D class averages. (**C-E**) Local resolution estimation (0.5 FSC criterion) and Fourier Shell Correlation (FSC) curves with global resolution (0.143 FSC criterion) for (C) the $(NIAUF)_2$ consensus refinement (EMD-19355), (D) the local refinement FDX2-bound proximal (EMD-19356), and (E) the local refinement FDX2-bound distal (EMD-19357). (**F**) Viewing direction distribution plot for the C2 consensus refinement.

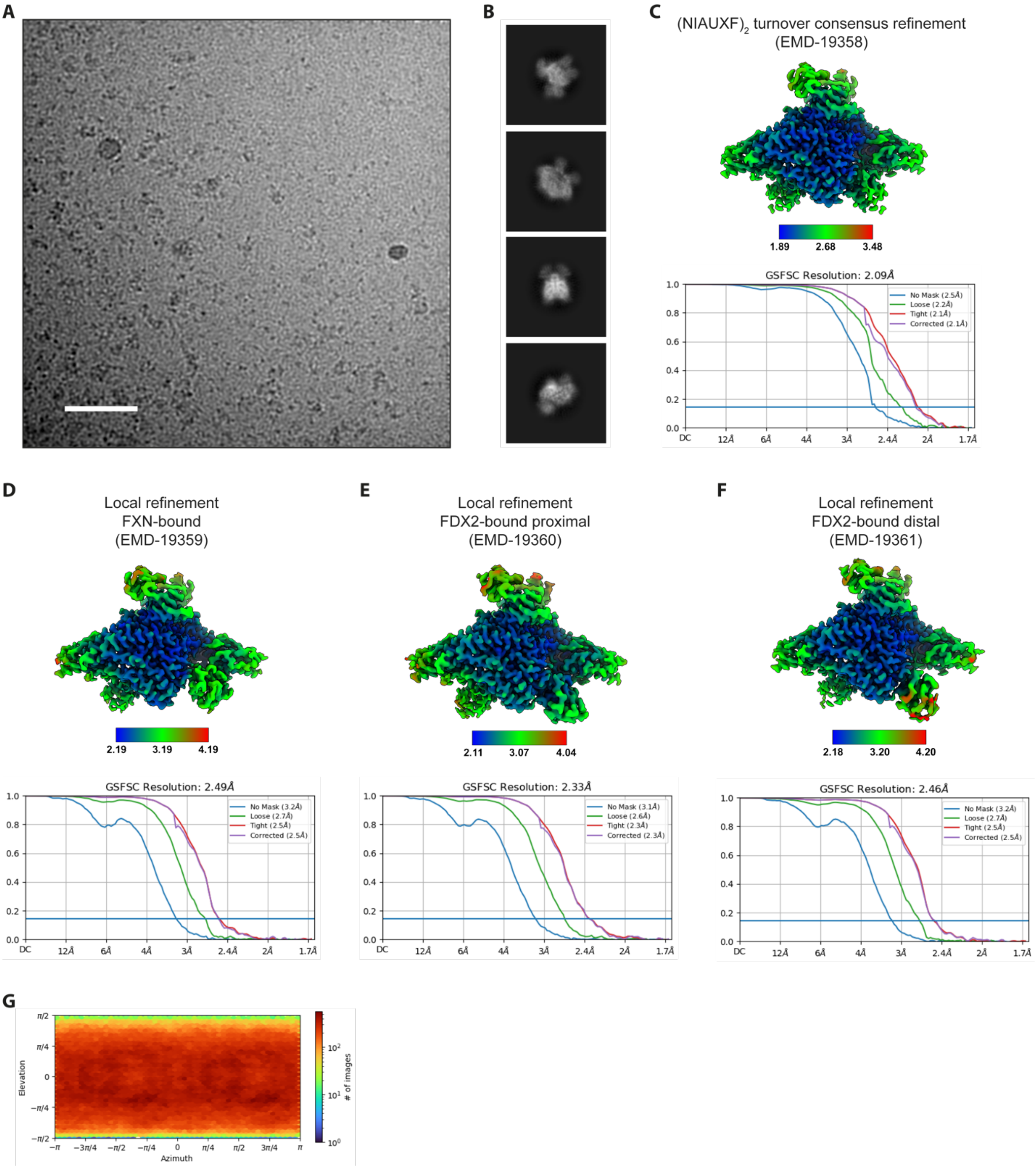


**Fig. S2. Cryo-EM data processing of the $(NIAUXF)_2$ turnover dataset.**

(**A**) Representative cryo-EM micrograph (scale bar 40 nm), (**B**) representative 2D class averages, **(C-F)** Local resolution estimation (0.5 FSC criterion) and Fourier Shell Correlation (FSC) curves with global resolution (0.143 FSC criterion) for (C) the $(NIAUXF)_2$ turnover consensus refinement (EMD-19358), (D) local refinement FXN-bound (EMD-19359), (E) local refinement FDX2-bound proximal (EMD-19360), (F) local refinement FDX2-bound distal (EMD-19361). (**G**) Viewing direction distribution plot for the C2 consensus refinement.

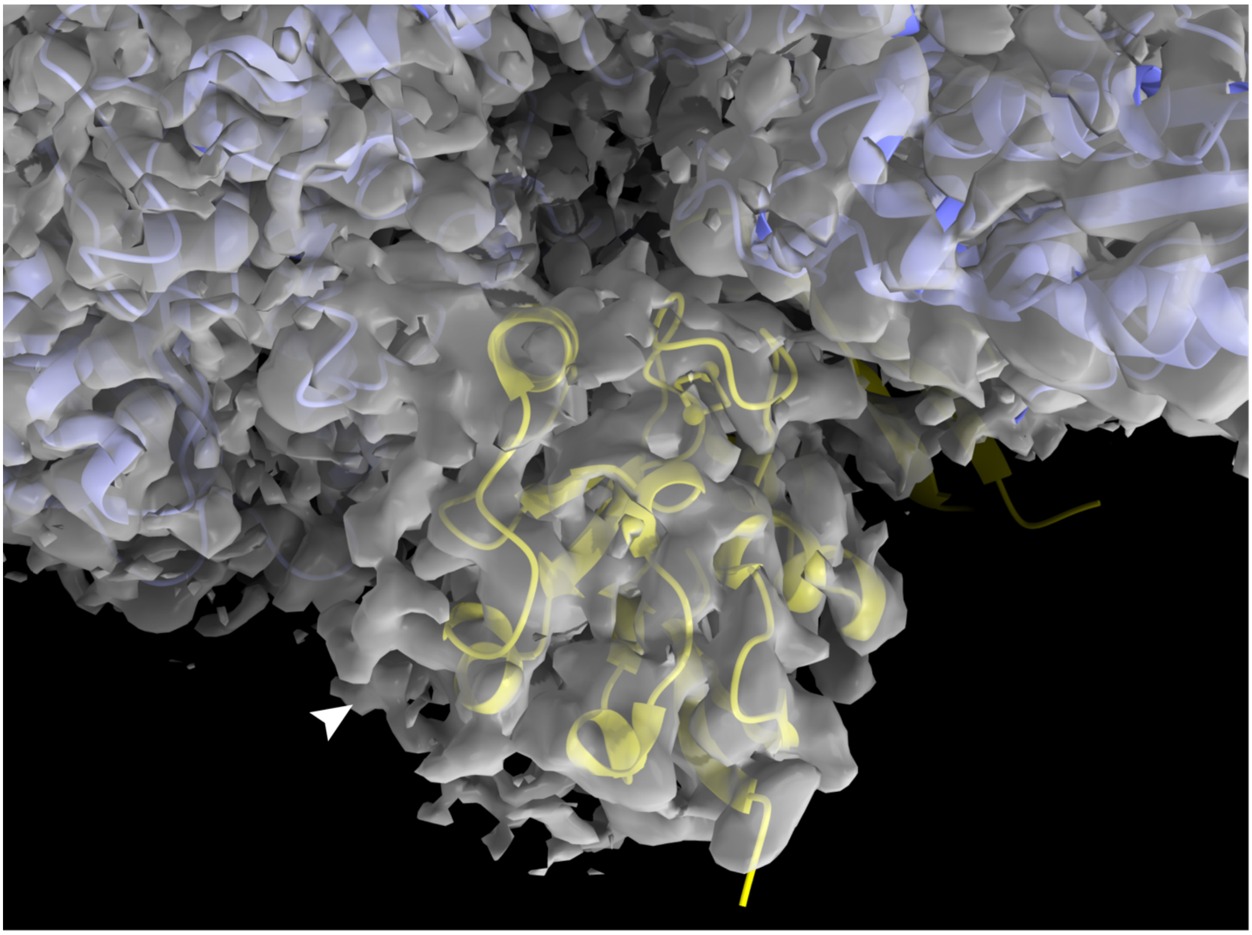

**Fig. S3. FDX2-binding region in the $(NIAUF)_2$ consensus refinement map.**
The atomic model of the FXN-bound core ISC complex ($(NIAUX)_2$, PDB 6NZU; purple), from which FXN was deleted, and the model of FDX2 (PDB 2Y5C; lemon) were rigid-body fitted into the consensus refinement map. Unmodelled density in the map (arrow) is due to heterogeneity in the position of FDX2 binding.

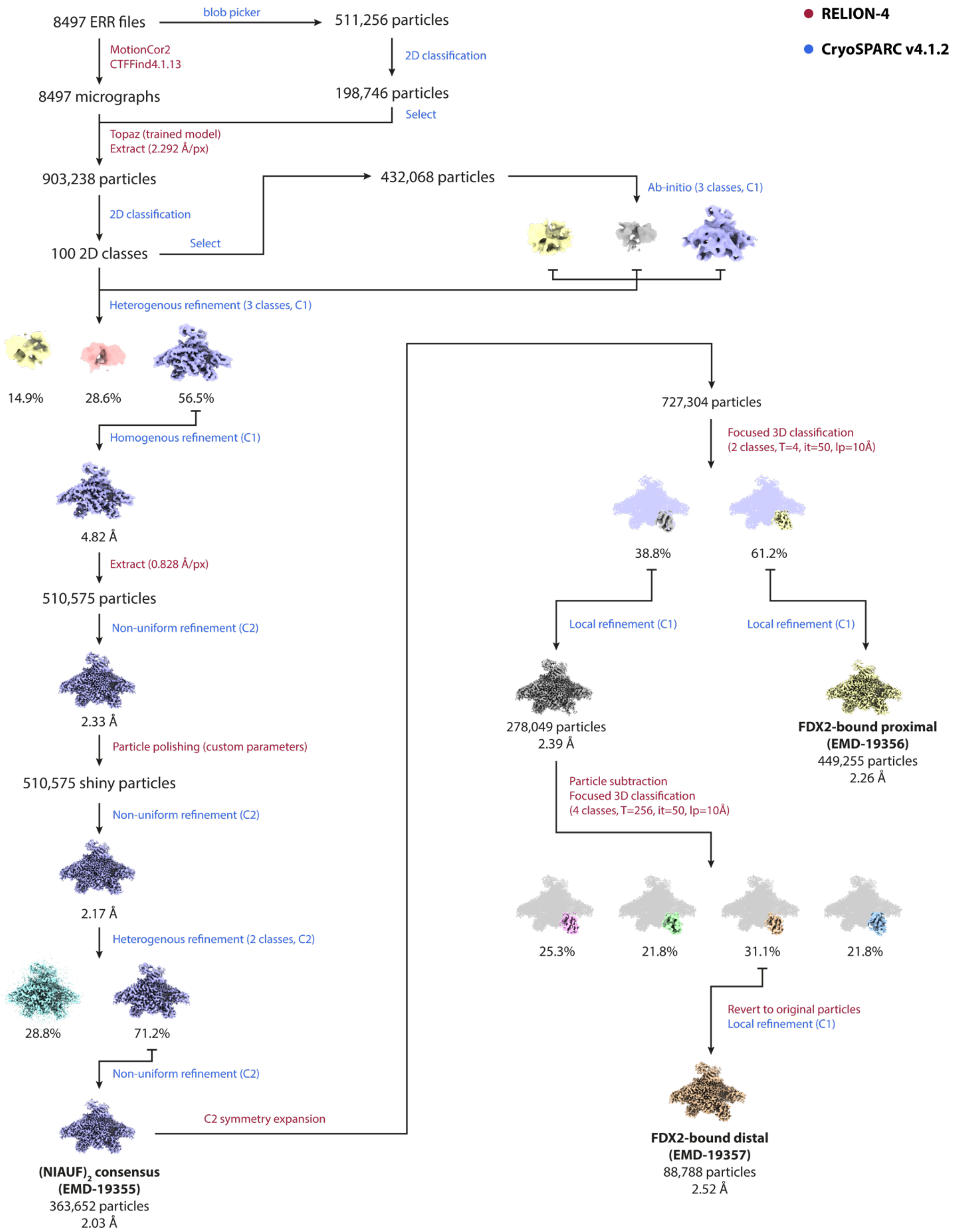


**Fig. S4. Data processing scheme for the (NIAUF)$_2$ dataset.**
Processing steps were performed in RELION-4 (red) and CryoSPARC v4.1.2 (blue).

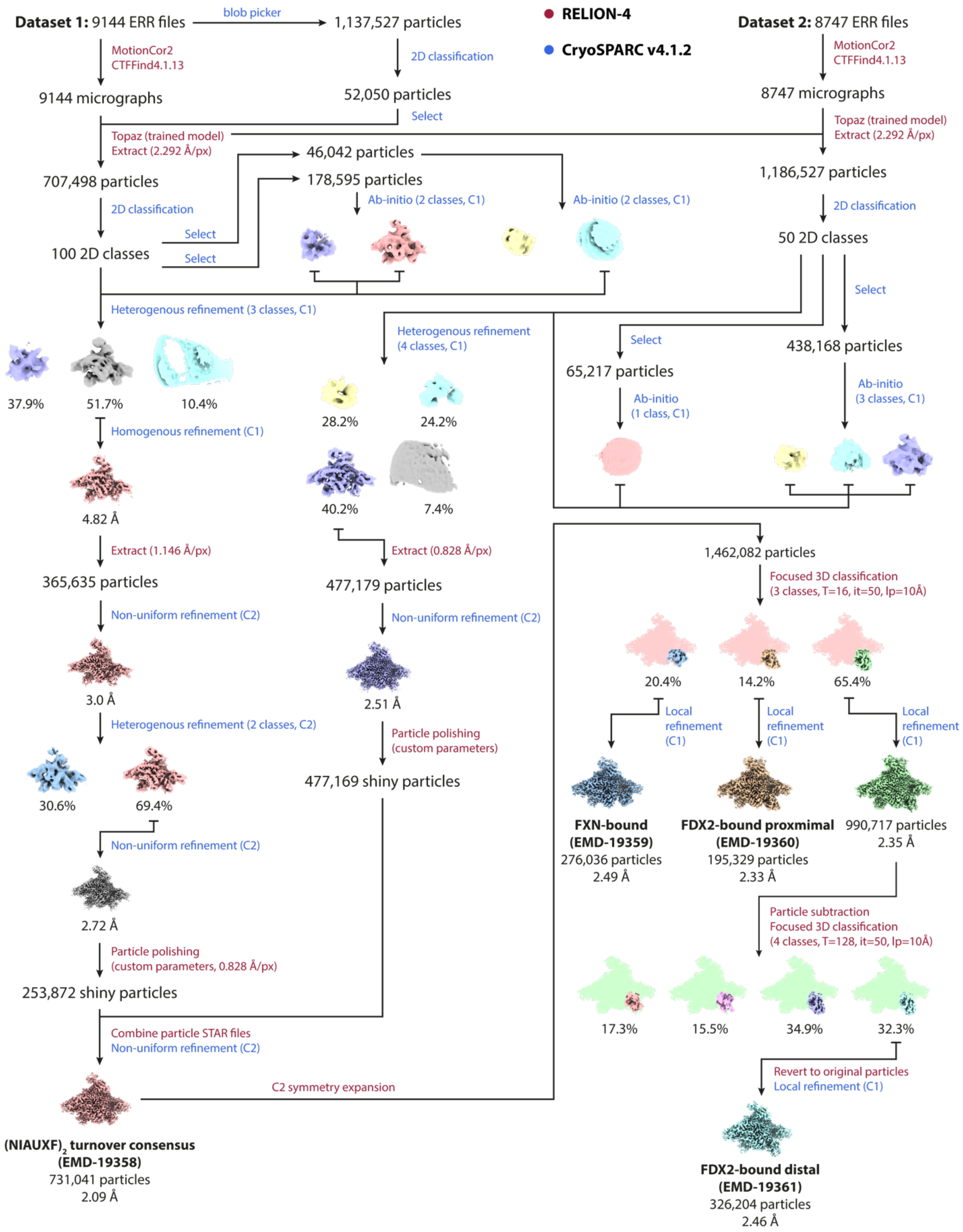


**Fig. S5. Data processing scheme for the (NIAUXF)$_2$ turnover datasets.**
Processing steps were performed in RELION-4 (red) and CryoSPARC v4.1.2 (blue).

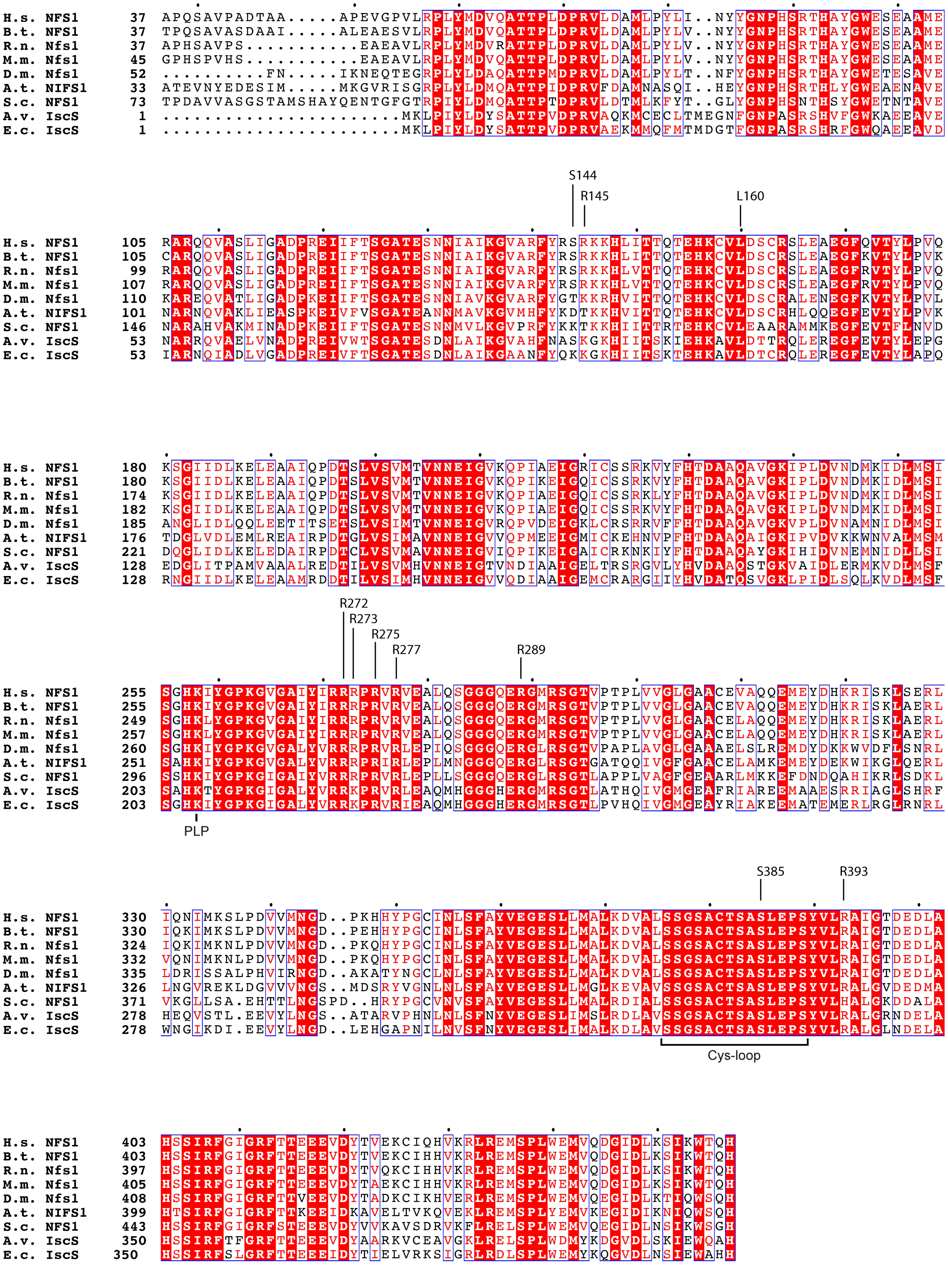
S144
R145
L160
R272
R273
R275
R277
R289
PLP
S385
R393
Cys-loop

**Fig. S6. Multi-sequence alignment of NFS1-like proteins.**

The Cys-loop region, the PLP cofactor linked to Lys258 and residues interacting with human FDX2 are annotated. Sequence identifiers: *H. sapiens* NFS1 (Q9Y697), *B. taurus* NFS1 (A5PKG4), *R. norvegicus* Nfs1 (Q99P39), *M. musculus* Nfs1 (Q9Z1J3), *D. melanogaster* Nfs1 (Q9VKD3), *A. thaliana* NIFS1 (O49543), *S. cerevisiae* NFS1 (P25374), *A. vinelandii* IscS (O31269), *E. coli* IscS (P0A6B7).

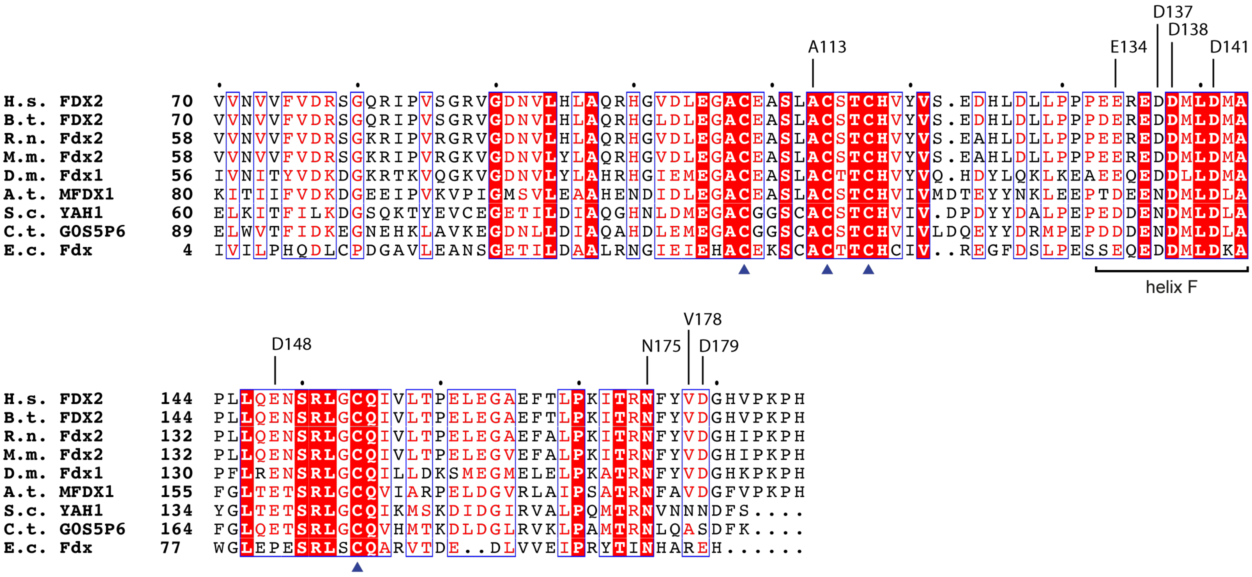


**Fig. S7. Multi-sequence alignment of FDX2-like proteins.**

Residues interacting with NFS1/NFS1’ are annotated. Blue triangles indicate the [2Fe-2S] cluster-coordinating cysteines Cys108, Cys114, Cys117 and Cys154 of human FDX2. Sequence identifiers: *H. sapiens* FDX2 (Q6P4F2), *B. taurus* FDX2 (Q05B51), *R. norvegicus* Fdx2 (D4A8N2), *M. musculus* Fdx2 (Q9CPW2), *D. melanogaster* Fdx1 (P37193), *A. thaliana* MFDX1 (Q9M0V0), *S. cerevisiae* YAH1 (Q12184), *C. thermophilum* Putative 2 iron, 2 sulfur cluster binding protein (G0S5P6), *E. coli* Fdx (P0A9R4).

**Table S1. Protein sequences.**

Brackets indicate sequences that were removed by proteolytic cleavage and are not present in the final sample preparation.

| Protein | Sequence | Uniprot ID |
|---|---|---|
| NFS1 | MSLRPLYMDVQATTPLDPRVLDAMLPYLINYYGNPHSRTHAYGWESEAAMERARQQ<br>VASLIGADPREIIFTSGATESNNIAIKGVARFYRSRKKHLITTQTEHKCVLDSCRSLEAEG<br>FQVTYLPVQKSGIIDLKELEAAIQPDTSLVSVMTVNNEIGVKQPIAEIGRICSSRKVYFHT<br>DAAQAVGKIPLDVNDMKIDLMSISGHKIYGPKGVGAIYIRRRPRVRVEALQSGGGQER<br>GMRSGTVPTPLVVGLGAACEVAQQEMEYDHKRISKLSERLIQNIMKSLPDVVMNGDP<br>KHHYPGCINLSFAYVEGESLLMALKDVALSSGSACTSASLEPSYVLRAIGTDEDLAHSSI<br>RFGIGRFTTEEEVDYTVEKCIQHVKRLREMSPLWEMVQDGIDLKSIKWTQH | Q9Y697-1 |
| ISD11 | MGSSHHHHHHGSPTTENLYFQGHNMAASSRAQVLALYRAMLRESKRFSAYNYRTYA<br>VRRIRDAFRENKNVKDPVEIQTLVNKAKRDLGVIRRQVHIGQLYSTDKLIIENRDMPRT | Q9HD34 |
| ACP1 | MGSDMPPLTLEGIQDRVLYVLKLYDKIDPEKLSVNSHFMKDLGLDSLDQVEIIMAMED<br>EFGFEIPDIDAEKLMCPQEIVDYIADKKDVYE | O14561 |
| ISCU2 | MAYHKKVVDHYENPRNVGSLDKTSKNVGTGLVGAPACGDVMKLQIQVDEKGKIVDA<br>RFKTFGCGSAIASSSLATEWVKGKTVEEALTIKNTDIAKELCLPPVKLHCSMLAEDAIK<br>AALADYKLKQEPKKGEAEKKLEHHHHHH | Q9H1K1-1 |
| FXN | (MHHHHHHSSGVDLGTENLYFQ)SNASGTLGHPGSLDETTYERLAEETLDSLAEFFEDL<br>ADKPYTFEDYDVSFGSGVLTVKLGGDLGTYVINKQTPNKQIWLSSPSSGPKRYDWTGK<br>NWVYSHDGVSLHELLAAELTKALKTKLDLSSLAYSGKDA | Q16595 |
| FDX2 | MASDVVNVVFVDRSGQRIPVSGRVGDNVLHLAQRHGVDLEGACEASLACSTCHVYVS<br>EDHLDLLPPPEEREDDMLDMAPLLQENSRLGCQIVLTPELEGAEFTLPKITRNFYVDGH<br>VPKPH | Q6P4F2 |
| FDXR | MGSSHHHHHHSQDPNSTQEKTPQICVVGSGPAGFYTAQHLLKHPQAHVDIYEKQPVPF<br>GLVRFGVAPDHPEVKNVINTFTQTAHSGRCAFWGNVEVGRDVTVPELREAYHAVVLS<br>YGAEDHRALEIPGEELPGVCSARAFVGWYNGLPENQELEPDLSCDTAVILGQGNVALD<br>VARILLTPPEHLERTDITKAALGVLRQSRVKTVWLVGRRGPLQVAFTIKELREMIQLPG<br>ARPILDPVDFLGLQDKIKEVPRPRKRLTELLLRTATEKPGPAEAARQASASRAWGLRFF<br>RSPQQVLPSPDGRRAAGVRLAVTRLEGVDEATRAVPTGDMEDLPCGLVLSSIGYKSRP<br>VDPSVPFDSKLGVIPNVEGRVMDVPGLYCSGWVKRGPTGVIATTMTDSFLTGQMLLQ<br>DLKAGLLPSGPRPGYAAIQALLSSRGVRPVSFSDWEKLDAEEVARGQGTGKPREKLVD<br>PQEMLRLLGH | P22570 |

**Table S2. Plasmids used for protein expression**

| Plasmid | ORF | Reference |
|---|---|---|
| pASK-IBA43(+)-*FDX2* | *FDX2 (1-68Δ)* | (Webert *et al.*, 2011) |
| pET24b(+)-*ISCU2* | *ISCU2-His*$_6$ *(1-34Δ)* | (Freibert *et al.*, 2021) |
| pETDuet1-*NFS1-ISD11* | *NFS1 (1-55Δ), His*$_6$*-Tev-ISD11* | (Freibert *et al.*, 2021) |
| pRSFDuet1-*ACP* | *ACP (1-68Δ)* | (Freibert *et al.*, 2021) |
| pMCSG7-*FXN* | *His*$_6$*-Tev-FXN (1-80Δ)* | (Freibert *et al.*, 2021) |
| pETDuet1-*FDXR* | *His*$_6$*-FDXR (1-32Δ)* | (Sheftel *et al.*, 2010) |

### Table S3. Data collection parameters, processing parameters and model validation statistics for the $(NIAUF)_2$ dataset

| | $(NIAUF)_2$ consensus map | $(NIAUF)_2$ FDX2-bound proximal | $(NIAUF)_2$ FDX2-bound distal |
|---|---|---|---|
| ***Map EMDB ID*** | **EMD-19355** | **EMD-19356** | **EMD-19357** |
| **Data collection** | | | |
| Microscope | Krios G4 | Krios G4 | Krios G4 |
| Camera | Falcon 4 | Falcon 4 | Falcon 4 |
| Voltage (kV) | 300 | 300 | 300 |
| Nominal magnification | 215,000x | 215,000x | 215,000x |
| Calibrated pixel size (Å) | 0.573 | 0.573 | 0.573 |
| Dose ($e^-$/Å$^2$) | 80 | 80 | 80 |
| Number of frames per image | 952 | 952 | 952 |
| Defocus range (µm) | -2.2 - -0.8 | -2.2 - -0.8 | -2.2 - -0.8 |
| **Image processing** | | | |
| Motion correction software | MotionCor2 | MotionCor2 | MotionCor2 |
| CTF estimation software | CTFFIND4 | CTFFIND4 | CTFFIND4 |
| Particle selection software | Topaz | Topaz | Topaz |
| Final micrographs (no.) | 8497 | 8497 | 8497 |
| Initial particle images (no.) | 903,238 | 903,238 | 903,238 |
| Final particle images (no.) | 363,652 | 449,255 (symmetry expanded) | 88,788 (symmetry expanded) |
| Symmetry applied | C2 | C1 | C1 |
| Map sharpening B-factor (Å$^2$) | -58.2 | * | * |
| Final resolution (Å) | 2.03 | 2.26 | 2.39 |
| *FSC threshold* | 0.143 | 0.143 | 0.143 |
| ***Model PDB ID*** | | **8RMC** | **8RMD** |
| **Refinement** | | | |
| Modeling software | | Coot, PHENIX | Coot, PHENIX |
| Protein residues | | 1482 | 1461 |
| Water | | 400 | 290 |
| Ligands | | 8Q1, FE2, FES, PLP | 8Q1, FE2, FES, PLP |
| **Validation** | | | |
| MolProbity score | | 1.35 | 1.48 |
| Clash score | | 4.72 | 6.25 |
| Ramachandran plot (%) | | | |
| Outliers | | 0.00 | 0.00 |
| Allowed | | 1.50 | 1.66 |
| Favored | | 98.50 | 98.34 |
| Rotamer outliers (%) | | 1.40 | 1.51 |
| Cβ outliers (%) | | 0.07 | 0.00 |
| Peptide plane (%) | | | |
| Cis proline/general | | 3.3/0.0 | 3.5/0.0 |
| Twisted proline/general | | 0.0/0.0 | 0.0/0.0 |
| CaBLAM outliers (%) | | 0.55 | 0.70 |

* the primary map was density modified with *phenix.resolve_cryo_em*

**Table S4. Data collection parameters, processing parameters and model validation statistics for the $(NIAUXF)_2$ turnover datasets.**

| | **$(NIAUXF)_2$ turnover, consensus map** | **$(NIAUXF)_2$ turnover, FXN-bound** | **$(NIAUXF)_2$ turnover, FDX2-bound proximal** | **$(NIAUXF)_2$ turnover, FDX2-bound distal** |
|---|---|---|---|---|
| ***Map EMDB ID*** | **EMD-19358** | **EMD-19359** | **EMD-19360** | **EMD-19361** |
| **Data collection** | | | | |
| Microscope | Krios G4 | Krios G4 | Krios G4 | Krios G4 |
| Camera | Falcon 4 | Falcon 4 | Falcon 4 | Falcon 4 |
| Voltage (kV) | 300 | 300 | 300 | 300 |
| Nominal magnification | 215,000x | 215,000x | 215,000x | 215,000x |
| Calibrated pixel size (Å) | 0.573 | 0.573 | 0.573 | 0.573 |
| Dose ($e^-/Å^2$) | 80 | 80 | 80 | 80 |
| Number of frames per image | 987 /1,078 | 987 / 1,078 | 987 / 1,078 | 987 / 1,078 |
| Defocus range (µm) | -2.2 - -0.8 | -2.2 - -0.8 | -2.2 - -0.8 | -2.2 - -0.8 |
| **Image processing** | | | | |
| Motion correction software | MotionCor2 | MotionCor2 | MotionCor2 | MotionCor2 |
| CTF estimation software | CTFFIND4 | CTFFIND4 | CTFFIND4 | CTFFIND4 |
| Particle selection software | Topaz | Topaz | Topaz | Topaz |
| Final micrographs (no.) | 17,891 | 17,891 | 17,891 | 17,891 |
| Initial particle images (no.) | 1,894,025 | 1,894,025 | 1,894,025 | 1,894,025 |
| Final particle images (no.) | 731,041 | 276,036 (symmetry expanded) | 195,329 (symmetry expanded) | 326,204 (symmetry expanded) |
| Symmetry applied | C2 | C1 | C1 | C1 |
| Map sharpening B-Factor ($Å^2$) | -68.2 | -60 | -58.2 | -69.1 |
| Final resolution (Å) | 2.09 | 2.49 | 2.33 | 2.46 |
| *FSC threshold* | 0.143 | 0.143 | 0.143 | 0.143 |
| ***Model PDB ID*** | | **8RME** | **8RMF** | **8RMG** |
| **Refinement** | | | | |
| Modeling software | | Coot, PHENIX | Coot, PHENIX | Coot, PHENIX |
| Protein residues | | 1466 | 1466 | 1463 |
| Water | | 157 | 253 | 158 |
| Ligands | | 8Q1, FE2, PLP | 8Q1, FE2, FES, PLP | 8Q1, FE2, FES, PLP |
| **Validation** | | | | |
| MolProbity score | | 1.54 | 1.35 | 1.53 |
| Clash score | | 5.41 | 5.74 | 6.32 |
| Ramachandran plot (%) | | | | |
| Outliers | | 0.00 | 0.00 | 0.00 |
| Allowed | | 2.35 | 2.14 | 1.52 |
| Favored | | 97.65 | 97.86 | 98.48 |
| Rotamer outliers (%) | | 1.67 | 0.87 | 1.75 |
| Cβ outliers (%) | | 0.00 | 0.07 | 0.15 |
| Peptide plane (%) | | | | |
| Cis proline/general | | 3.7/0.0 | 3.4/0.0 | 3.5/0.0 |
| Twisted proline/general | | 0.0/0.0 | 0.0/0.0 | 0.0/0.0 |
| CaBLAM outliers (%) | | 0.49 | 0.77 | 0.70 |

**Movie S1**

Morph between the distal (PDB 8RMD) and proximal (PDB 8RMC) $(NIAUF)_2$ conformations, showing the rotation of FDX2 helix F along the NFS1 arginine patch and movement of the ISCU2 Cys69-loop. The FDX2 C terminus, which is only resolved in the proximal conformation, was omitted for simplicity.
